# Tissue-specific metabolic reprogramming during wound induced *de novo* organ formation in tomato hypocotyl explants

**DOI:** 10.1101/2021.04.29.441912

**Authors:** Eduardo Larriba, Ana Belén Sánchez García, Cristina Martínez-Andújar, Alfonso Albacete, José Manuel Pérez-Pérez

**Affiliations:** Instituto de Bioingeniería, Universidad Miguel Hernández, 03202 Elche, Spain; CEBAS-CSIC, Department of Plant Nutrition, Campus Universitario de Espinardo, 30100 Espinardo, Murcia, Spain; CEBAS-CSIC, Department of Plant Nutrition, Campus Universitario de Espinardo, 30100 Espinardo, Murcia, Spain; current address: Institute for Agri-Food Research and Development of Murcia (IMIDA), Department of Plant Production and Agrotechnology, 30150 La Alberca, Murcia, Spain

**Author notes:** Corresponding author: José Manuel Pérez-Pérez.

**Keywords:** *de novo* root regeneration, *de novo* shoot apical meristem formation, time-course bulk RNA-Seq, photosynthesis, photorespiration, glycolysis/gluconeogenesis, reactive oxygen species (ROS)

## Abstract

Plants have remarkable regenerative capacity, which allows them to survive tissue damaging after biotic and abiotic stress. Some of the key transcription factors and the hormone crosstalk involved in wound-induced organ regeneration have been extensively studied in the model plant *Arabidopsis thaliana*. However, little is known about the role of metabolism in wound-induced organ regeneration.
Here, we performed detailed transcriptome analysis and targeted metabolomics approach during *de novo* organ formation in tomato hypocotyl explants and found tissue-specific metabolic differences and divergent developmental pathways after wounding.
Our results indicate that callus growth in the apical region of the hypocotyl depends on a specific metabolic switch involving the upregulation of the photorespiratory pathway and the differential regulation of photosynthesis-related genes and of the gluconeogenesis pathway.
The endogenous pattern of ROS accumulation in the apical and basal region of the hypocotyl during the time-course were dynamically regulated, and contributed to tissue-specific wound-induced regeneration.
Our findings provide a useful resource for further investigation on the molecular mechanisms involved in wound-induced organ formation in a crop species such as tomato.

**One-sentence Summary:** Metabolic switch during wound-induced regeneration

## INTRODUCTION

Plants have remarkable regenerative capacity, enabling them to either repair damaged tissues or regenerate a lost organ (Ikeuchi et al., 2019). Plants regenerative capacity allows them to overcome tissue damage caused by different environmental insults. Moreover, enhanced plant regeneration capabilities have been frequently used during crop production to increase yield by grafting and pruning without killing the plant, as well as for the clonal propagation of elite genotypes.

The many advantages of *Arabidopsis thaliana* and the wide repertoire of regenerative responses in this species have provided some insight into the molecular mechanisms underlying tissue repair and organ regeneration (Ikeuchi et al., 2016; Ikeuchi et al., 2020). Two broad categories of regenerative models have been established, hereafter referred to as tissue culture-induced regeneration and wound-induced regeneration (Mathew and Prasad, 2021). Key transcription factor families and the hormone crosstalk involved in plant regeneration have been extensively studied in this species (Ibáñez et al., 2020). In metazoans, transcription-factor mediated reprogramming of somatic cells into induced pluripotent stem cells (Takahashi and Yamanaka, 2006) involves a profound metabolic shift from oxidative phosphorylation to glycolytic-dependent energy generation (Panopoulos et al., 2012; Cliff and Dalton, 2017). In addition, rewiring of energetic metabolism in cancer cells has been proven essential for tumor progression (Frezza, 2020). Strikingly, in plant tissue culture-induced regeneration, addition of sugars, amino acids and polyamines to the growing media often influence cellular proliferation and regeneration efficiency, although the precise mechanism by which they act is currently unknown. Ectopic expression of Arabidopsis *WOUND INDUCED DEDIFFERENTIATION1* (*WIND1*) in hypocotyl explants of *Brassica napus* enhanced callus formation and root regeneration, a phenotype that correlated with deregulation of metabolites, such as sucrose, proline, gamma aminobutyric acid or putrescine (Iwase et al., 2018). However, the precise link between the activation of a single transcription factor with non-hormone metabolite accumulation and callus formation and regeneration requires additional investigations.

The ‘Micro-Tom’ cultivar of tomato (*Solanum lycopersicum* L.) has been recognized as a model for tomato research because it shares some key advantages with Arabidopsis including its small size, short life cycle and easy transformation capability (Shikata and Ezura, 2016). High-efficient targeted gene editing using an optimized CRISPR-Cas9 system (Dahan-Meir et al., 2018) and TILLING collections (Okabe et al., 2013) facilitates the use of reverse genetics approaches. In our previous studies on wound-induced adventitious root (AR) formation in tomato (Alaguero-Cordovilla et al., 2020; Alaguero-Cordovilla et al., 2021), we noticed formation of *de novo* shoot meristems at the cut surface of the hypocotyl after removal of the shoot apex. Here, we performed detailed transcriptome analyses and targeted metabolomics approach during *de novo* organ formation (i.e., adventitious shoots and ARs) in tomato hypocotyl explants, to provide a conceptual framework for the identification of the regulatory mechanisms involved in organ regeneration. The findings of this study reveal the role of metabolism on tissue-specific wound-induced organ formation in tomato.

## MATERIALS AND METHODS

### Plant material and growth conditions

Seedlings of the tomato cultivar ‘Micro-Tom’ were grown *in vitro* as described elsewhere (Alaguero-Cordovilla et al., 2021). Hypocotyl explants were obtained after removing the whole root system (2-3 mm above the hypocotyl-root junction) and the shoot apex (just below the cotyledons’ petiole) with a sharp scalpel, at the 100-101 growth stages (Feller et al., 1995), hereafter 0 days after excision (dae). Hypocotyl explants were transferred to 120 × 120 mm (length × width) Petri dishes containing 75 mL of standard growing medium (SGM) (Alaguero-Cordovilla et al., 2021), otherwise indicated (**Supplemental Table 1a**).

### RNA isolation, library construction and NGS sequencing

For each sample, three to four mm of the apical or the basal regions of the hypocotyl were collected at 0, 1, 4 and 8 dae (denoted as T0, T1, T4, and T8). Three biological replicates, each consisting of 12 apical or basal fragments respectively, were obtained, and immediately frozen in liquid N2. Total RNA from ~100 mg of powdered tissue that was kept at −80 °C was extracted using Spectrum^™^ Plant Total RNA Kit (Sigma-Aldrich, USA) and treated with DNAse I (ThermoFisher Scientific, USA). The RNA integrity was confirmed using a 2100 Bioanalyzer (Agilent Technologies, USA). Three sequencing libraries from basal tissues and two sequencing libraries form apical tissues at T0, T1, T4 and T8 were obtained with the TruSeq Stranded RNA Sample Preparation Kit v2 (Illumina, USA). NGS sequencing was carried out by Macrogen (Korea) on an Illumina HiSeq4000 in pair-end mode with 100 cycles of sequencing. Additionally, three NGS libraries were sequenced using the BGISEQ-500 pipeline at BGI Genomics (BGI Tech Solutions, Hong Kong, China), in pair-end mode with 150 cycles of sequencing. The raw NGS reads (SRA number) were pre-processed using Trimmomatic (Bolger et al., 2014) and with FastQC (https://www.bioinformatics.babraham.ac.uk/projects/fastqc/) for quality assessment.

### RNA-Seq analysis

The bioinformatics workflow used is shown in **Supplemental Figure 1a**. Briefly, clean RNA-Seq reads were mapped to *Solanum lycopersicum* L. genome build SL4.0 (Hosmani et al., 2019) using STAR 2.7 (Dobin et al., 2013). Assignation of reads to gene models (ITAG4.0 annotation) was performed with featureCounts from the Subread package (Liao et al., 2019). Identification of significant transcriptional changes was carried out based on Bayesian estimation of temporal regulation (false discovery rate [FDR] = 0.05) (Aryee et al., 2009) implemented in MeV software package (http://mev.tm4.org/). Read count normalization and differential gene expression analysis were carried out using DESeq2 integrated into Differential Expression and Pathway analysis (iDEP 9.1) web application (Ge et al., 2018). Statistics and read count normalization were indicated in **Supplemental Figure 1b-g**. Differential expressed genes (DEG) were filtered using an FDR < 0.01 and Log2fold change > < |1|. K-means clustering and principal component analysis were carried out using normalized counts on the iDEP 9.1 web application. Web-tool ShinyGO v0.61 (Ge et al., 2020) was used to perform Gene Ontology (GO) enrichment analysis. ITAG4.0 gene structural annotation was retrieved from Sol Genomics Network (SolGenomics; https://solgenomics.net/). Their putative *Arabidopsis thaliana* orthologs were identified using BioMart tool from the Emsembl Plants database (Bolser et al., 2017). Genes were assigned to KEGG metabolic pathways by Reciprocal Best Hit BLAST using GhostKOALA (https://www.kegg.jp/ghostkoala/) and SolGenomics BLAST against ITAG4.0. For heatmap and hierarchical clustering analyses, we used Morpheus (https://software.broadinstitute.org/morpheus) webtool.

### Gene expression analysis by real-time quantitative PCR

Total RNA from ~100 mg of powdered hypocotyl sections from 14 seedlings at T0, T1, T4 and T8 was extracted in triplicate using Spectrum Plant Total RNA Kit (Sigma–Aldrich, United States) and further processed as described elsewhere (Alaguero-Cordovilla et al., 2021). For real-time quantitative PCR (RT-qPCR), primers amplified 115-205 base pairs of the cDNA sequences of selected genes (**Supplemental Table 1b**). RT-qPCR was performed as previously described (Alaguero-Cordovilla et al., 2021).

### Macroscopic studies of wound-induced organogenesis

Hypocotyl explants were incubated for 10 to 21 days in SGM supplemented with different compounds (**Supplemental Table 1b**). In all these cases, ARs arising from the hypocotyl were visually scored and periodically annotated. AR emergence was estimated based on the day before the observation of the first AR. Shoot regeneration stages were also scored based on the morphological structures observed in the apical region of the hypocotyl. For photorespiration assays, we applied the treatments on 0.2% agarose droplets (25 μL) at the apical region of the hypocotyl explants; shoot regeneration stages were scored at 17 dae.

### Cell death and reactive oxygen species (ROS) assays

For microscopic examination of dying cells, hypocotyl explants at T0, T1, T4 and T8 were stained on hot acetic acid/trypan blue solution (0.5% w/v in 45% acetic acid) for 3 minutes. After incubation for 1 hour at room temperature, hypocotyls were cleared in chloral hydrate (80 g chloral hydrate in 30 mL distilled water). For DAB staining, whole hypocotyl explants atT0, T1, T4 and T8 dae were immersed on a 1% (w/v) solution of DAB in Tris-HCl buffer, pH 4.8. After 1 hour of vacuum infiltration, samples were kept in DAB solution overnight at room temperature. Then, explants were cleared in chloral hydrate solution. To confirm the specific accumulation of H_2_O_2_, before DAB staining, we incubated hypocotyl explants with 100 units·μl^-1^ catalase (Sigma-Aldrich) for 30 minutes. Thereafter, samples were processed as indicated above.

### Metabolite extraction and analysis

Three biological replicates, each consisting of eight apical or basal regions of the hypocotyl were collected at T0, T1, T4 and T8. Metabolites were extracted from frozen tissues and were analyzed as described previously by (Albacete et al., 2008) with some modifications. Fresh plant material (0.1 g) was homogenized in liquid nitrogen and incubated in 1 mL of cold (−20 °C) extraction mixture of methanol/water (80/20, v/v) for 30 min at 4 °C. Solids were separated by centrifugation (20,000 g, 15 min at 4 °C) and re-extracted for another 30 min at 4 °C with 1 mL of extraction solution. Pooled supernatants were passed through Sep-Pak Plus C18 cartridges to remove interfering lipids and some plant pigments. The supernatant was collected and evaporated under vacuum at 40 °C. The residue was dissolved in 0.2 mL methanol/water (20/80, v/v) solution using an ultrasonic bath. The dissolved samples were filtered through 13 mm diameter Millex filters with 0.22 μm pore size nylon membrane (Millipore, Bedford, MA, USA) and placed into opaque microcentrifuge tubes.

Ten μL of filtered extract were injected in a U-HPLC-MS system consisting of an Accela Series U-HPLC (ThermoFisher Scientific, Waltham, MA, USA) coupled to an Exactive mass spectrometer (ThermoFisher Scientific, Waltham, MA, USA) using a heated electrospray ionization (HESI) interface. Mass spectra were obtained using the Xcalibur software version 2.2 (ThermoFisher Scientific, Waltham, MA, USA). Metabolites of interest were identified by extracting the exact mass from the full scan chromatogram obtained in the negative mode and adjusting a mass tolerance of ≤ 1 ppm. The concentrations were semi-quantitatively determined from the extracted peaks using the calibration curves of analog compounds.

### Statistical analyses

The descriptive statistics were calculated by using the StatGraphics Centurion XV software (StatPoint Technologies, Inc. Warrenton, VA, USA) and SPSS 21.0.0 (SPSS Inc., Chicago, IL, USA) programs. Outliers were identified and excluded for posterior analyses as described elsewhere (Alaguero-Cordovilla et al., 2021). We performed multiple testing analyses using the ANOVA F-test or Fisher’s least significant difference (LSD) methods (p-value < 0.01, otherwise indicated). Non-parametric tests were used when necessary (i.e. AR emergence and AR number).

## RESULTS

### Time-course RNA-Seq analysis during wound-induced organ formation in tomato hypocotyl explants

We previously studied wound-induced adventitious root (AR) development in the basal region of Micro-Tom hypocotyl explants (Alaguero-Cordovilla et al., 2021). We observed formative divisions of the cambial cells facing the phloem one day after wounding. AR primordia emerged from specific domains within the basal region of the explants after 3-4 days, and 96.8% of the explants developed between two and four ARs at 8 days (**Figure 1a**) (Alaguero-Cordovilla et al., 2021). Intriguingly, hypocotyl explants obtained by sectioning shoot explants just below the cotyledons developed new functional shoots at their apical end after three weeks (**Figure 1b**, rs). We confirmed the absence of pre-formed shoot apical meristems (SAM) in these explants at T0 that quickly sealed the wound in the apical region (**Figure 1c**, stage 1). We found that homogeneous callus-like tissue formation developed from 4-5 days onwards (**Figure 1c**, stage 2). After 8 days, some regionalization within these calluses occurred and bud outgrowth was obvious (**Figure 1c**, stage 3), which will form new shoot apices afterwards (**Figure 1c**, stage 4).

**Figure 1.**
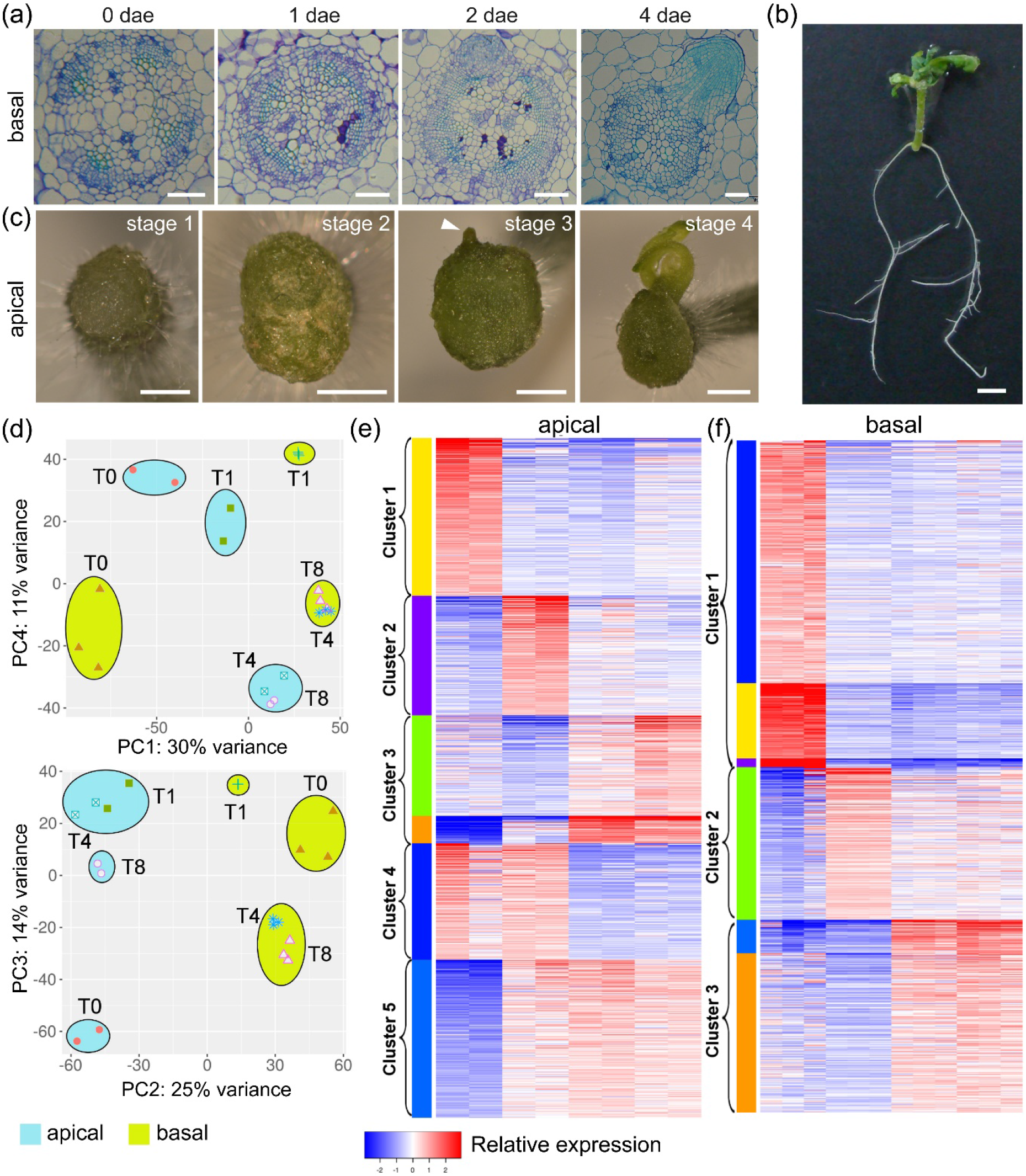
Wound-induced organ formation in tomato hypocotyl explants. (a) Adventitious root (AR) development in the basal region of the hypocotyl explants. (b) Whole plant regeneration from hypocotyl and stem node explants at 21 dae; rs: regenerated shoot. (c) Developmental stages during *de novo* shoot formation in the apical region of the hypocotyl. Scale bars: 100 μm (a), 1 mm (b) and 4 mm (c). (d) Principal component (PC) analysis of the RNA-Seq results. (e, f) K-means clustering of time-course RNA-Seq results from apical (e) and basal (f) regions of the hypocotyl. Predicted clusters are colored and grouped accordingly to its temporal expression profile.

To reveal the molecular signatures during *de novo* organ formation in tomato, we performed time-course bulk RNA-Seq analysis of apical and basal regions of the hypocotyl explants at 0, 1, 4 and 8 days after wounding (see **Materials and Methods**). We found that 20,831 genes displayed significant transcriptional changes, representing 60% of the tomato genes in ITAG4.0 annotation. About 10 % of transcripts were long non-coding RNA (lncRNA; **Supplemental Figure 1d, e)**. We assayed the expression profile of two genes by qRT-PCR (**Supplemental Figure 2a, b**), and these results supported the reliability of our RNA-Seq dataset. Principal component (PC) analysis of expression profiles showed differences in transcript profiling both between apical and basal regions and during the time-course (**Figure 1d**).

### Functional enrichment analysis of gene expression profile during wound-induced regeneration

K-means clustering analysis across different time points and hypocotyl regions, was performed using the most variable expressed genes (standard derivation, SD > 0.5), resulting in 4,500 and 4,600 genes from apical and basal regions, respectively (**Supplemental Figure 2c, d**). The optimal cluster number was estimated using the Elbow method implemented in iDEP.91 (Ge et al., 2018) (**Supplemental Figure 2e, f**). Three of these clusters displayed the same temporal profile in apical and basal regions, corresponding to genes mostly expressed at T0 (cluster 1), genes expressed at T1 (cluster 2), and genes expressed at T4 and T8 (cluster 3; **Figure 1e, f**). Besides, we found two additional clusters in the apical region, which included genes expressed at T0-T1 (cluster 4) and T1-T4-T8 (cluster 5; **Figure 1e**). Venn diagrams and Gene Ontology (GO) enrichment analysis in clusters 1-3 allowed us to identify either specific (apical or basal) or common GO enrichment subsets (**Supplemental Figure 3** and **Supplemental Table 2**).

Genes in cluster 1 might represent the basal transcriptome of young tomato hypocotyls and would reveal tissue-specific differences at T0. Enriched genes shared between apical and basal regions (10.6 %; **Supplemental Figure 3a** and **Supplemental Table 2**) were related to auxin signaling and gibberellin biosynthesis, which are known to contribute to hypocotyl growth (Jiang et al., 2020).

Cluster 2 might contain key regulators of *de novo* organ formation as their expression levels transiently increased 24 h after wounding (**Figure 1e, f**). We did not find significant GO enrichment for the common genes in this cluster (**Supplemental Table 2**), which was consistent with our PC analyses and with the divergent developmental patterns at the apical and basal regions after wounding (**Figure 1d)**. A GO enriched network for specific genes in the apical region in this cluster was related to nucleosome assembly and chromatin remodeling (**Figure 2a** and **Supplemental Table 2**). On the other hand, oxidation-reduction processes and GO terms related to hormone biosynthesis and regulation were among the mostly GO enriched terms found in the basal region (**Supplemental Table 2**). Genes in cluster 3 were mostly expressed between T4 and T8 (**Figure 1e, f**). The most-significantly enriched GO terms shared between apical and basal regions were related to cell cycle function (**Figure 2b** and **Supplemental Table 2**). Nevertheless, GO terms related to photosynthesis and *de novo* protein folding were highly enriched in the apical region (**Figure 2c** and **Supplemental Table 2**). On the other hand, highly enriched GO terms in the basal region belonged to cell wall modification and reactive oxygen species (ROS) detoxification (**Supplemental Table 2**). Cluster 4 genes were mostly expressed in the apical region up to 24 h after excision (T1) and were enriched in terms related to cell wall and polysaccharide metabolism (**Figure 1e** and **Supplemental Table 2**). Cluster 5 genes, which were highly expressed between T1 and T8 in the apical region, were enriched in GO terms related to carbon metabolism-related biosynthetic process (**Figure 1e** and **Supplemental Table 2**).

**Figure 2.**
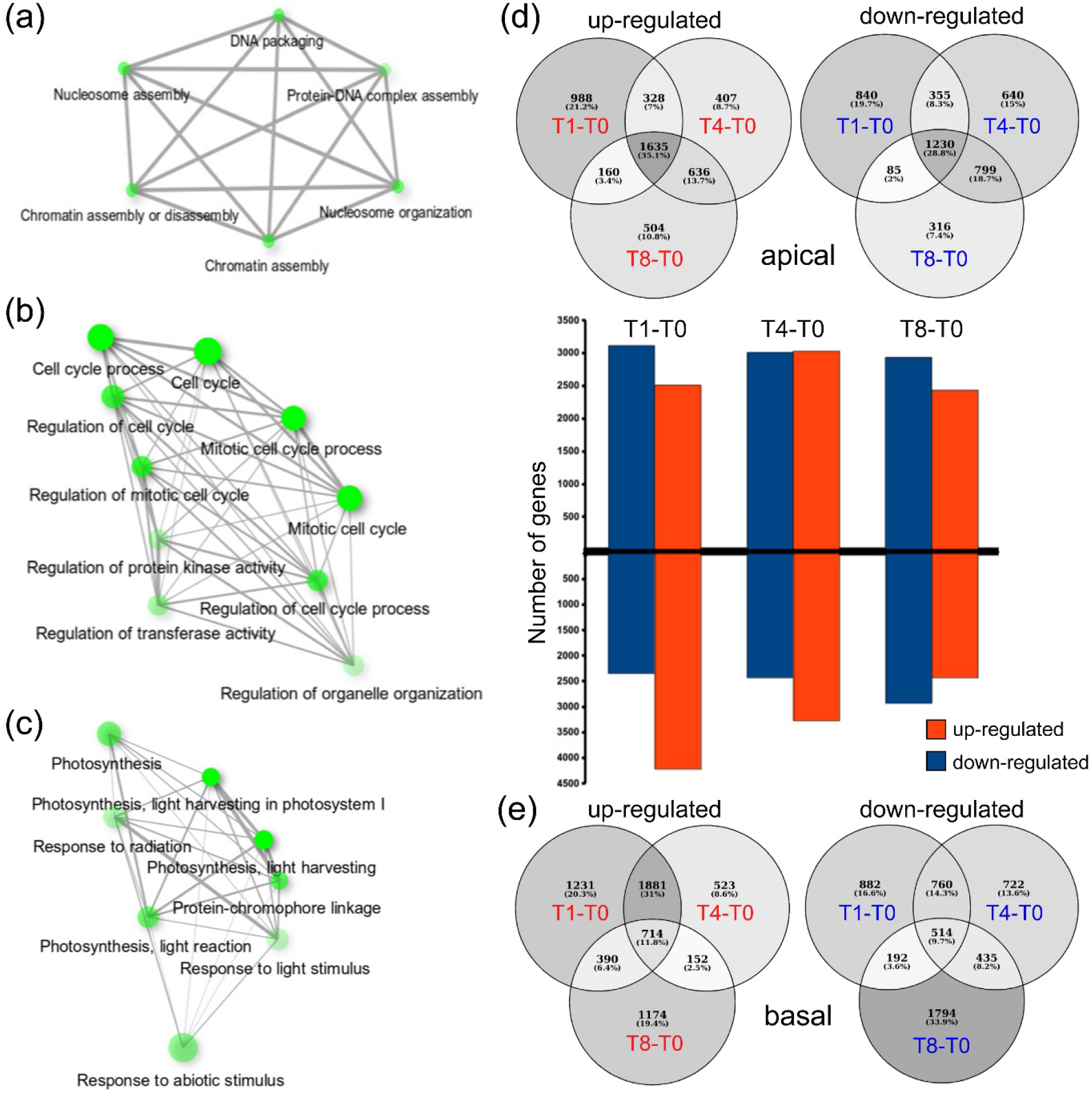
Functional enrichment and DEG analysis during wound-induced regeneration. (a-c) Examples of GO term-enriched networks of the upregulated genes in the apical region at T1 (a), of the upregulated genes at T4-T8 shared between apical and basal regions (b), and of specific upregulated genes in the apical region at T4-T8 (c). (d, e) Venn diagrams of upregulated and downregulated DEGs, from apical (d) and basal (e) regions of the hypocotyl as regards T0.

Our GO-enrichment analysis based on K-means gene clustering suggests that both shared and specific biological processes are taking place in the apical and basal regions of the explants during *de novo* organ formation.

### Differentially expressed genes during wound-induced regeneration

We identified 12,406 deregulated genes (DEG) as regards T0, representing 59.6% of the expressed genes in the RNA-Seq analysis. In the apical region of the explants, we found 8,792 DEG (4,658 and 4,265 upregulated and downregulated genes, respectively), whose overlap in different contrasts (**Figure 2d**) allowed us to perform GO term enrichment analysis of specific gene subsets. Within the upregulated genes from T1 to T8 in the apical region, we found enrichment of GO terms associated with serine/glycine metabolism, sterol biosynthesis, and response to oxidative stress (**Supplemental Table 3**). Besides, some genes related to hormone response were downregulated and significantly enriched through the time-course in this region (**Supplemental Table 3**). Downregulated genes at T1 were enriched in GO terms related to photosynthesis (light-dependent reactions) and *de novo* protein folding, while a subset of auxin response genes were upregulated in T1 (**Supplemental Table 3**). Interestingly, we found upregulated genes that were enriched in GO terms related to cell cycle regulation, some of which were shared between T1 and T8 contrasts, while others were specific of T8 (**Supplemental Table 3**). Within the downregulated genes at the T4 and T8 contrasts, we found GO terms related to cell wall biogenesis and cell redox homeostasis (**Supplemental Table 3**). Overlapping with leaf primordia formation in this region, some genes annotated as being involved in leaf development were also significantly upregulated at T8 (**Supplemental Table 3**).

We found 9,085 DEG (4,076 and 5,233 upregulated and downregulated genes, respectively) in the basal region as regards T0 (**Figure 2e**). Within the upregulated genes from T1 to T8 (**Supplemental Table 3**), we found enrichment of GO terms associated with ion transport. Some photosynthesis-related genes were downregulated at T1, whereas other genes were also downregulated through the time-course (**Supplemental Table 3**). The GO terms related to protein translation, rRNA modification and RNA processing were enriched at the T1 contrast (**Supplemental Table 3**). For DEG related to cell cycle, a subset of genes involved in cell division was downregulated at T1 while other genes were upregulated at T4 (**Supplemental Table 3**). We also found a contrasting regulation of genes assigned to GO terms related to response to oxidative stress. Indeed, some genes involved in ROS detoxification (mainly encoding peroxidase function) were up-regulated at T4 and T8 contrasts, while genes involved in H_2_O_2_ responses were down-regulated from T1 to T8 (**Supplemental Table 3**). The GO term enrichment for upregulated genes at T4 and T8 highlighted genes involved in cell wall biogenesis, which follows the formation and emergence of AR primordia in the basal region.

The biological functions identified by GO term enrichment analyses might contribute to the separate developmental functional processes of tissue reprogramming during *de novo* shoot formation and AR formation in the apical and basal regions of the tomato hypocotyl explants, respectively, and thus deserve further attention.

### Tissue-specific regulation of photosynthesis during wound-induced organ formation

Our functional GO analysis showed an enrichment in genes related to photosynthesis process during *de novo* organ formation (**Supplemental Table 2** and **3**). Due to the relevance of this metabolic process, we have identified 90 genes in the photosynthesis-related KEGG pathway databasetime-course (**Supplemental Table 4**), 76 of them (84.4%) were deregulated (**Supplemental Figure 4a, b**). Hierarchical clustering analysis of gene expression values highlighted two predominant profiles in the apical region: genes mainly repressed at T1, but that they regain their expression at T4 and T8 (red cluster, **Figure 3a, c),** and genes with an increase in expression from T1 to T8 (purple cluster, **Figure 3a, c**). On the other hand, most photosynthesis-related genes in the basal region showed strong repression in T1 and mild decreased expression at T4 and T8 as regards T0 levels (green cluster, **Figure 3b, c**). These results showed differential gene expression regulation of the photosynthetic machinery in the apical and basal regions after wounding (**Figure 3d-g**). Our DEG analysis uncovered genes involved in regulating the plastoquinone pool (**Supplemental Figure 4a, c**), such as those encoding the NADPH subunits PNSB1 and PNSB4, or those encoding ferredoxins (FD) and the NADP^+^ reductases LFNR1 and PGR5-like, which were upregulated in the apical region from T1 onwards (**Figure 3d**). In the apical region, most genes encoding core components of PSII were upregulated from T1 onwards (**Figure 3e**). However, most of the genes encoding different PSI proteins were downregulated at T1 in this region, and showed a strong upregulation at T4 and T8 in the apical region (**Figure 3f**). Interestingly, genes encoding antenna proteins of both photosystems (LHCB and LHCA) displayed a severe downregulation of their expression levels in the apical region only at T1 and were downregulated in the basal region from T0 onwards (**Supplemental Figure 4b, d**). In addition to deregulation of genes associated with different components of the photosynthetic machinery in the apical region, regulatory genes encoding the protein kinases ABC1K1 and STN8, as well as the auxiliary PSII core protein PSB33, were also upregulated (**Figure 3g**). Conversely, in the basal region all these genes were significantly downregulated from T1 onwards (**Figure 3g**), suggesting that photosynthesis function might be reduced in the basal region after wounding and during AR formation.

**Figure 3.**
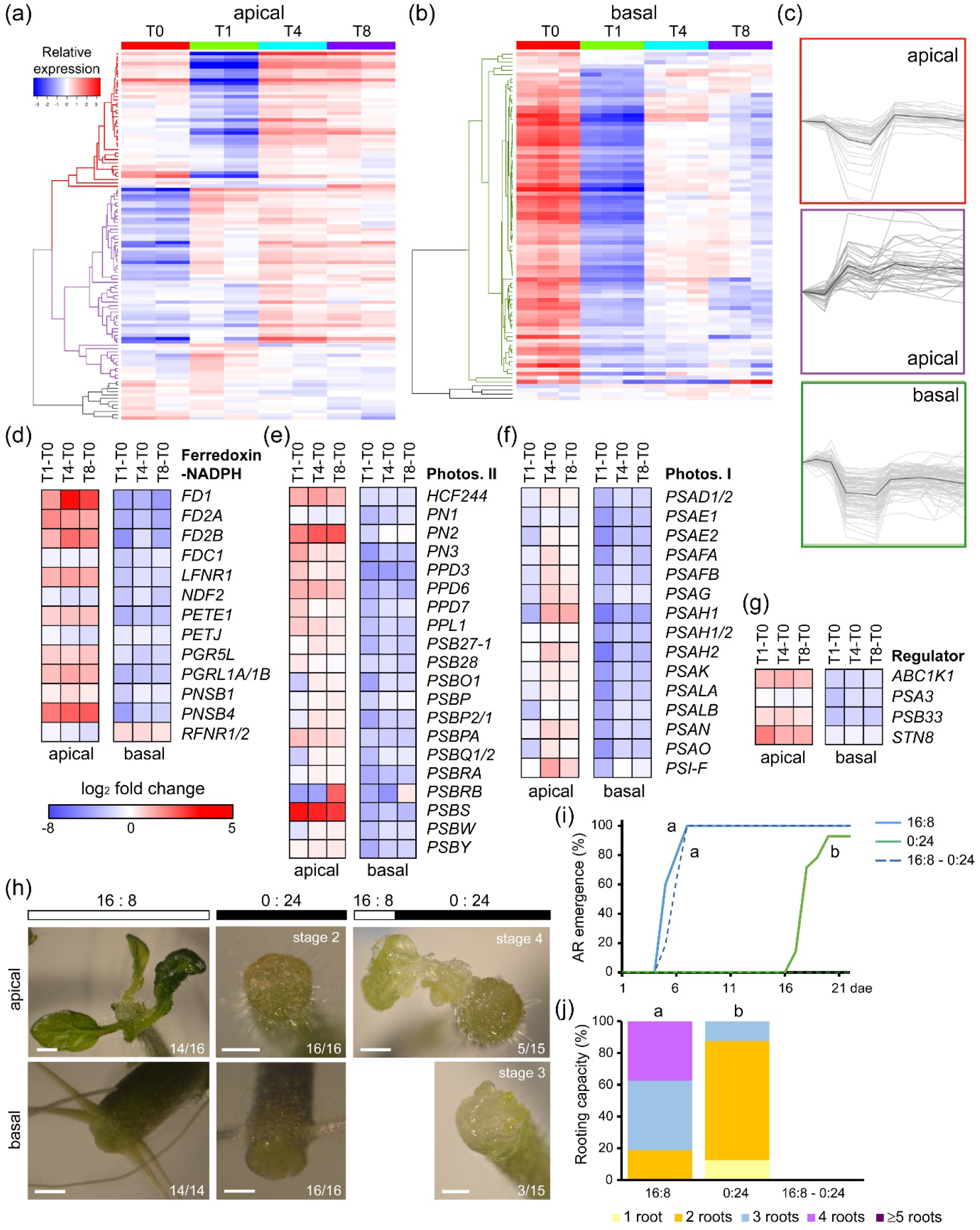
Regulation of photosynthesis-related genes during wound-induced organ formation. (a, b) Hierarchical clustering of the time-course expression of photosynthesis genes in the apical (a) and basal (b) regions of the hypocotyl explants. (c) Expression profiles of some of the gene clusters identified. (d-g) DEG involved in know photosynthesis subcomplexes, such as (d) ferredoxin:NAPDH reductase, (e) photosystem II or (f) photosystem I, and in (g) key regulators. (h) Representative images of wound-induced organ formation in the apical and basal region of the hypocotyl explants on different photoperiod conditions: 21-days standard photoperiod (16 h light and 8 h darkness; 16:8), 21-days darkness (0 h light and 24 h darkness; 0:24), and 3 days on 16:8 photoperiod followed by 18 days on 0:24 photoperiod. Gene annotations (d-g) are found in Supplemental Table 4. Scale bars: 1 mm.

To determine the photosynthesis requirement for wound-induced organ formation, we grown hypocotyl explants on different light and photoperiod conditions (**Figure 3h-l**). Hypocotyl explants grown on continuous darkness (0:24 photoperiod) were able to produce ARs in the basal region of the hypocotyl, albeit with severe delay in their emergence (**Figure 3 h, i**), but *de novo* organ formation in the apical region of these explants was held at stage 2 (**Figure 3 h**). We found that hypocotyl explants incubated for 3 days on standard photoperiod conditions (16 h light and 8 h darkness; 16:8) and then transferred to continuous darkness, were able to overcome stage 2 and did produce new shoots (**Figure 3h**). Taken together, these results indicate that wound-induced *de novo* shoot formation in hypocotyl explants requires the light signal.

### Tight regulation of ROS homeostasis is required for wound-induced AR formation

Because of photooxidative stress after wounding, light-induced ROS accumulation might occur in hypocotyl explants. Indeed, we found GO term enrichment of genes related to ROS detoxification during the studied time-course (**Supplemental Table 2** and **3)**. To measure ROS production, we visualized H_2_O_2_ accumulation by DAB staining during wound-induced organ formation (see **Materials and Methods**). At T1, higher DAB staining was found in the apical region of the explants (**Figure 4a**). Interestingly, the DAB precipitate was restricted to the tissue near the wound at T4, both in the apical and the basal region of the explants (**Figure 4a**, T4 inset). At T8, DAB staining was much lower and restricted to the most distal cells in the basal region and specific cells within the apical callus (**Figure 4a**, T8 inset). We confirmed DAB-specific staining of H_2_O_2_ accumulation by catalase incubation (**Figure 4a**, T1 cat).

**Figure 4.**
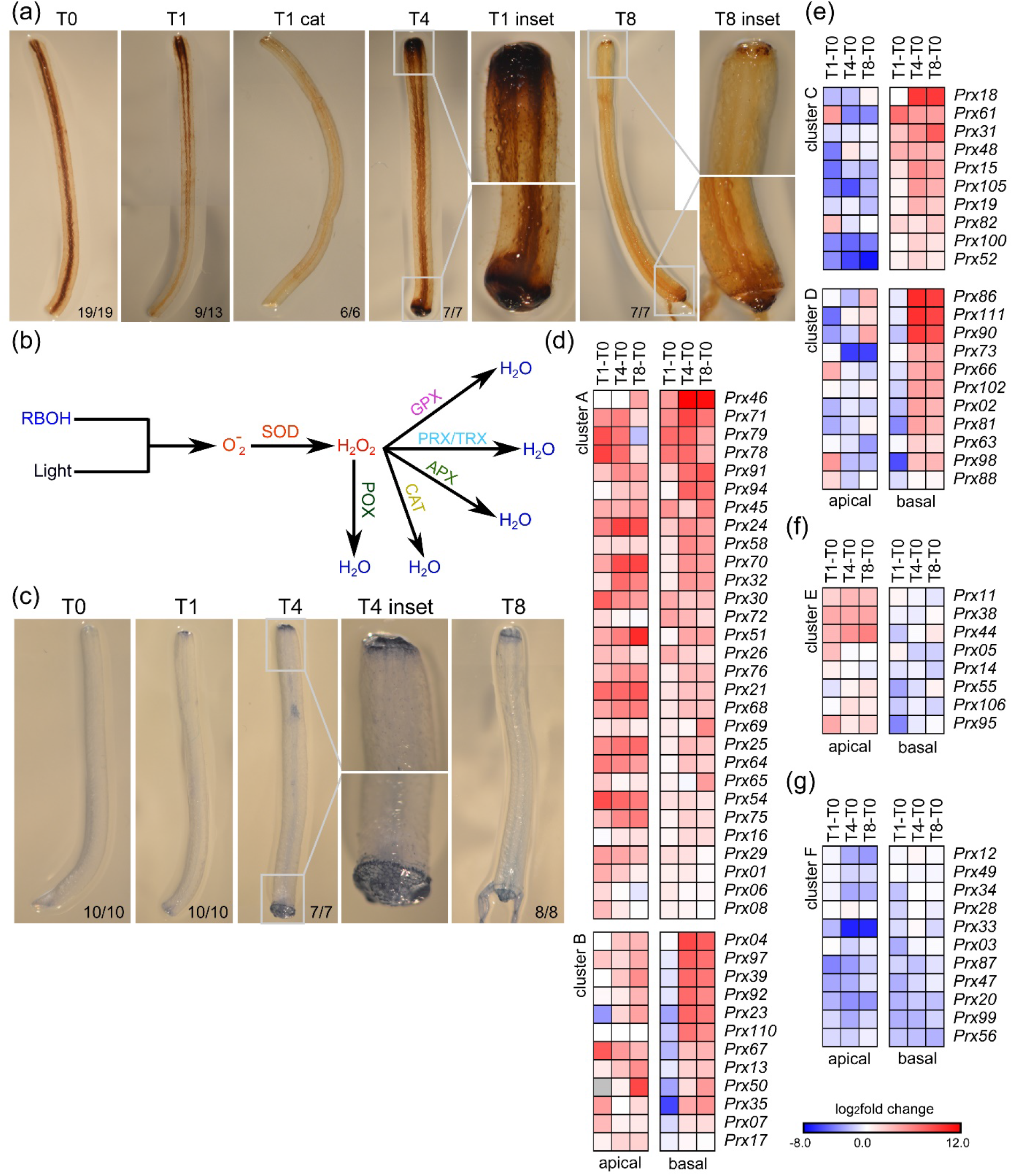
Reactive oxygen species (ROS) regulation during wound-induced organ formation. (a) H_2_O_2_ accumulation visualized by DAB staining in the hypocotyl explants during the studied time-course. (b) Proposed pathways of ROS production and detoxification with key enzymatic activities indicated: RBOH, NADPH oxidase/respiratory burst oxidase homolog; SOD, superoxide dismutase; POX, peroxidase; GPX, glutathione peroxidase; PRX/TRX, peroxiredoxins/thioredoxins; APX, ascorbate peroxidases; CAT, catalases. (c) Localized cell death in the hypocotyl explants during the studied time-course visualized by trypan blue staining. (d-f) Hierarchical clustering of DEG encoding class III peroxidases. Gene annotations are found in Supplemental Table 5. Scale bars (a, c): 4 mm.

To get additional insight into ROS regulation, we retrieved 135 expressed genes related to ROS homeostasis (**Supplemental Table 5**) that were assigned to different families based on the enzymatic activities of their products (Savelli et al., 2019). A hundred and ten (81.4%) of these genes were DEG (**Figure 4b**). In the apical region, most of the genes encoding ROS scavenging proteins were upregulated as regards T0 (**Supplemental Figure 5a, b**). In addition, five genes encoding membrane-dependent NADPH oxidases of the RBOH family (**Supplemental Figure 5a, b**), involved in stress-induced singlet oxygen production (Sagi et al., 2004; Devireddy et al., 2021), were upregulated both in the apical and the basal region (**Supplemental Figure 5b**). As excessive singlet oxygen might cause increased ROS levels that eventually lead to cell death (Gill and Tuteja, 2010), we visualized cell death by trypan blue staining (see **Materials and Methods**; **Figure 4c**). At T1, we found a faint trypan blue staining in the distal end of the apical region and no trypan blue staining in the basal region (**Figure 4c**, T1). At T4, however, trypan blue stained the outer cell layer of the basal region and the most distal cell layer of the apical region of the explants (**Figure 4c**, T4 inset). Similar trypan blue staining was found for distal cells within the apical and the basal regions of the hypocotyl explants at T8 **(Figure 4c**, T8), indicating that the extent of ROS was not directly associated with cell death. Genes encoding peroxiredoxins (PRX) and thioredoxins (TRX), ascorbate peroxidases (APX), and catalases (CAT) were differentially regulated between the apical and the basal regions (**Supplemental Figure 5a, b**), which might account for the observed differences in H_2_O_2_ accumulation upon wounding (**Figure 4a**).

We gathered 85 expressed genes encoding class III peroxidases (POX), 81 of which (95.3%) were deregulated largely than other ROS scavenging proteins (**Supplemental Table 5**). We grouped them accordingly to six different expression profiles (**Figure 4d-g**). Forty-one DEG were constitutively upregulated in both the apical region and the basal region during the time-course (e.g. *Prx54*, *Prx68*), with a few of these genes (e.g. *Prx92*, *Prx97*) being downregulated at T1 in the basal region (clusters A and B, respectively; **Figure 4d**). Twenty-nine DEG showed contrasting deregulation in the apical and the basal regions, with 21 of them being upregulated mainly in the basal region (e.g. *Prx100*, *Prx105*, clusters C and D; **Figure 4e**), and eight DEG were upregulated in the apical region only (e.g. *Prx44*, *Prx95*, cluster E; **Figure 4f**). Eleven DEG were mostly downregulated in both tissues as regards T0 (e.g. *Prx20*, *Prx87*, cluster F; **Figure 4g**). Our results indicate differential regulation of POX-encoding genes in the apical and the basal region of the hypocotyl explants during the time-course, which might account for dynamic ROS homeostasis during wound-induced organ formation in tomato hypocotyl explants.

### Photorespiration is required for wound-induced *de novo* shoot formation

We found 315 expressed genes related to carbon metabolism of which 199 were DEG (**Supplemental Table 6**). Key metabolic pathways, such as glycolysis/gluconeogenesis, carbon fixation through the Calvin-Benson cycle, glycolate/glyoxylate metabolism (i.e. photorespiration) and pentose phosphate pathway, were deregulated in our bulk time-course RNA-Seq (**Supplemental Figure 6a, b**). A subset of these DEG showed contrasting deregulation between apical and basal regions and deserved further attention. Because of the massive downregulation of the photosynthesis machinery at T1 (see above), the energy supply required for new organ growth might be compromised on the hypocotyl explants. Deregulation of genes encoding key enzymes of the Calvin-Benson cycle for the CO_2_ fixation via 3-phosphoglycerate (3-PGA), showed opposite trends in the apical and basal regions as regards T0, which suggest *de novo* establishment of sink-source relationships after wounding in these two regions (**Supplemental Figure 6b**).

Photorespiratory metabolism involves the production of 3-PGA by the oxygenase activity of the RuBisCO enzyme through the glycine and serine pathway at a high metabolic cost (Dellero et al., 2016). Because most genes encoding RuBisCO subunits are highly expressed (**Figure 5a**), and the atmospheric O_2_ floods the wounded tissue in the apical region, photorespiration should be active in this tissue. We found that several genes encoding key enzymes of the glycine biosynthesis pathway downstream of 2-phosphoglycolate, such as PGLP1, GLO4, and GGAT, were specifically upregulated as regards T0 in the apical region (**Figure 5a**). In contrast, all these genes were downregulated in the basal region after wounding (**Figure 5a**). Because of this differential regulation, glycine levels, but also H_2_O_2_ as a sub-product, might increase largely in the apical region. In the next step of the photorespiratory pathway, serine is produced via the glycine decarboxylase complex (GDC) and serine hydroxymethyltransferase (SHM) enzymes (Dellero et al., 2016). *GLP1/2*, which encodes the P-subunit of the GDC, and *SHM1* were upregulated in the apical region as regards T0 (**Figure 5a**). Serine is converted to hydroxypyruvate by the product of *AGT1*, which was also found upregulated in the apical region (**Figure 5a**). Finally, hydroxypyruvate is incorporated into the Calvin-Benson cycle as 3-PGA through the enzyme activities encoded by *HPR* and *GLYK* (**Figure 5a**). As it was found for the previous steps, most of the other genes of the photorespiratory pathway were not deregulated in the basal region after wounding, which clearly indicates that photorespiration is restricted to the apical region during wound-induced organ regeneration in tomato hypocotyl explants. To confirm that the observed gene expression changes contribute to a spatial regulation of photorespiration, we measured the endogenous levels of three pathway metabolic intermediaries (glycolate, glyoxylate and hydroxypyruvate) during *de novo* organ formation by targeted metabolome analysis (see **Materials and Methods**). At wounding time (T0), we did not find significant differences in endogenous levels of the studied intermediaries between apical and basal regions of the explants (**Figure 5b**). However, in all cases we found highly significant differences in the endogenous levels of these intermediaries during the time-course, with the highest levels found in the apical region between T1 and T8 (**Figure 5b**). In agreement with these results, 3-PGA levels were significantly enhanced in the apical region of the explants at T1 and T4, while there were no significant differences at T8 between apical and basal regions (**Figure 5c**). 3-PGA is a key intermediary of both the Calvin-Benson cycle and the glycolysis, which is converted into glyceraldehyde 3-phosphate (G3P) by the phosphoglycerate kinase (PGK) and the glyceraldehyde-phosphate dehydrogenase (GAPA) (**Figure 5a**). We found higher levels of 1,3-bisphosphoglycerate and glyceraldehyde 3-phosphate in the apical region of the explants (**Figure 5c**), which were in agreement with a higher upregulation of the *PGK* and *GAPA* genes in this region (**Figure 5a**). In addition, the genes encoding the enzymes involved in the following steps for regeneration of ribulose 1,5-bisphosphatase in the Calvin-Benson cycle were also differentially regulated in the apical and the basal regions (**Supplemental Figure 6b**).

**Figure 5.**
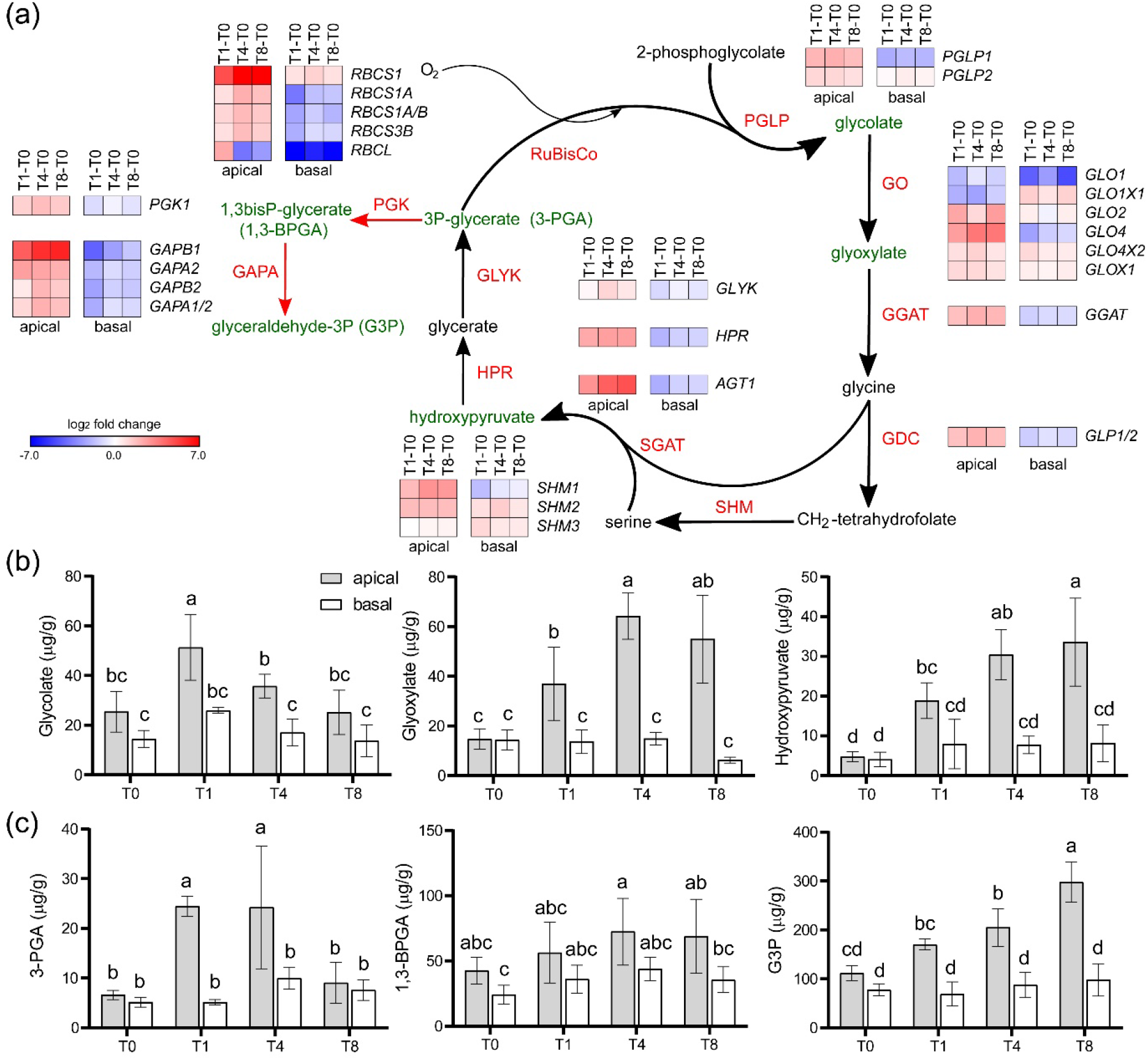
Photorespiration pathway activation during wound-induced organ formation. (a) DEG involved in photorespiration. Genes encoding the enzymatic activities of the different steps in the pathway are shown: PGLP, phosphoglycolate phosphatase; GO, glycolate oxidase; GGAT, glutamate-glyoxylate aminotransferase; GDC, glycine decarboxylase complex; SHM, serine hydroxymethyltransferase; SGAT, serine-glyoxylate aminotransferase; HPR, hydroxypyruvate reductase; GLYK, glycerate kinase. Two enzymes of the Calvin-Benson cycle are also included: PGK, phosphoglycerate kinase; GAPA, glyceraldehyde-phosphate dehydrogenase. (b, c) Endogenous levels of several metabolites of the photorespiration pathway in the apical and basal region of the hypocotyl explants during the studied time-course. Different letters indicate significant differences (p-value<0.01) between the assayed conditions. Gene annotations (a) are found in Supplemental Table 6.

Taken together, these results indicate that the apical region suffers a strong metabolic reprogramming after wounding, being photorespiration a restricted pivotal metabolic pathway that is required for effective *de novo* shoot formation after wounding.

### Spatial regulation of sugar metabolism during wound-induced organ formation

Due to the presence of sucrose in the regeneration medium, the explants might metabolize sugars through the glycolysis pathway as a direct energy source. Invertases irreversibly hydrolyze sucrose into glucose and fructose (Wan et al., 2018), and are classified accordingly to their subcellular localization in cell wall, cytosolic and vacuolar invertases (CWIN, CIN and VIN; **Supplemental Table 7**) (Proels and Hückelhoven, 2014; Tauzin and Giardina, 2014). Besides, sucrose cleavage is mediated by cytosolic sucrose synthase (SUS) (Stein and Granot, 2019). Most genes encoding CIN, VIN and SUS enzymes were downregulated as regards T0 in the apical and basal regions (**Figure 6a**). On the other hand, several *CWIN* genes were upregulated in both regions as regards T0 expression (**Figure 6a**). Extracellular glucose and fructose are taken up by plant cells via hexose transporters (Chen et al., 2015; Julius et al., 2017) (**Supplemental Table 7**). We found that several genes encoding SWEET transporters (Manck-Götzenberger and Requena, 2016) were upregulated in apical and basal regions of the explants (e.g. *SWEET10b*, *10c*, *11a*, *12a*, *12b* and *12c*) during the time-course (**Figure 6b**). Additionally, *SUT1*, whose product is involved in apoplastic phloem loading (Slewinski et al., 2009), was specifically upregulated in the apical region during the time-course (**Figure 6b**).

**Figure 6.**
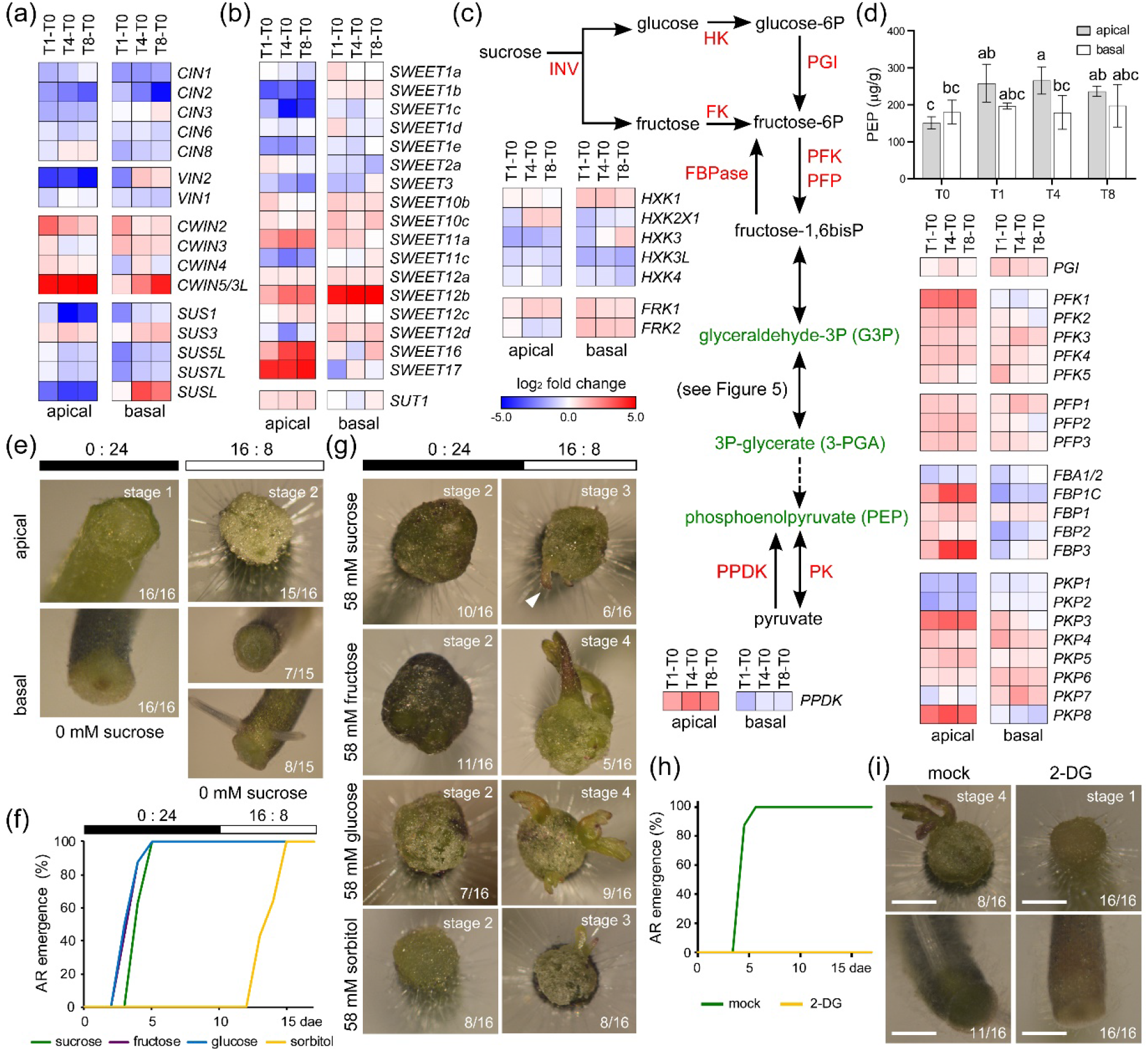
Sucrose is required for wound-induced organ formation. (a) DEG encoding invertases (CIN, VIN and CWIN) and sucrose synthases (SUS). (b) DEG encoding putative sugar transporters of the SWEET and SUT families. (c) Some DEG encoding enzymes in the glycolysis/gluconeogenesis pathway. Genes encoding the key enzymatic activities of the different steps in the pathway are shown: INV, invertase; HK, hexokinase; FK, fructokinase; PGI, phosphoglucose isomerase; PFK/PFP, phosphofructokinase; FBPase, fructose bisphosphatase; PK, pyruvate kinase; PPDK, pyruvate-phosphate dikinase. (d) Endogenous levels of phosphoenolpyruvate in the apical and basal region of the hypocotyl explants during the studied time-course. (e) Representative images of *de novo* organ formation in the apical and basal region of the hypocotyl explants on different photoperiod conditions (16:8 and 0:24) without sucrose. (f) AR emergence of hypocotyl explants grown for 10 days on 0:24 photoperiod followed by 7 days on 16:8 photoperiod. (g) Representative images of shoot formation stages in the apical region of the hypocotyl explants grown on different sugars with same photoperiod conditions as in (f). (h) Rooting capacity of hypocotyl explants at 14 dae on SGM (mock and 2-DG) and 16:8 photoperiod. Different letters (d, f, h) indicate significant differences (p-value<0.01) between the assayed conditions. (i) Representative images of *de novo* organ formation in the apical and the basal region of the hypocotyl explants on SGM (mock and 2-DG) and 16:8 photoperiod. Gene annotations (a-c) are found in Supplemental Tables 6 and 7. Scale bars (e, g, i): 1 mm.

Metabolic activation of glucose and fructose involves hexokinase (HK) and fructokinase (FK) enzymes, respectively, which were expressed and were differentially regulated in our RNA-seq (**Figure 6c** and **Supplemental Table 7**). The first downstream regulatory step of the glycolysis pathway involves ATP-dependent phosphofructokinase (PFK) and the pyrophosphate-dependent phosphofructokinase (PFP). We identified eight genes encoding PFK enzymes and five genes encoding PFP enzymes (**Supplemental Table 7)**, most of which were upregulated in both regions during the time-course (**Figure 6c**). Several genes encoding pyruvate kinase (PK), another key regulatory enzyme of the glycolytic pathway, were similarly upregulated in the apical and basal regions as regards T0 (**Figure 6c**). These results suggest that sucrose is catabolized to pyruvate both in the apical and the basal regions, while photorespiration provides a surplus of G3P that might also be used for pyruvate production mostly in the apical region (see previous section). Indeed, pyruvate levels slightly increased in the apical region but were not significantly affected in the basal region during the time-course (**Figure 6d**). Importantly, gluconeogenesis might also occur from pyruvate or oxaloacetate, the latter being a direct product of the citrate cycle or the glycolate/glyoxylate cycle. Indeed, the gene encoding the pyruvate, phosphate dikinase (PPDK) was found upregulated in the apical region of the explants during the time-course (**Figure 6c**). Another key regulatory factor in gluconeogenesis is fructose 1,6-bisphosphatase (FBPase), and we found three FBPase-encoding genes that were specifically upregulated in the apical region of the explants from T1 onwards (**Figure 6c**). Taken together, these results confirm the metabolic switch of the apical region from being a sink to becoming a source tissue.

### *De novo* organ formation in tomato hypocotyl explants depends on sugar availability

Finally, since we identified that both photosynthesis and sucrose metabolism are deregulated in wound-induced organ formation, we assayed the relevance of these two factors during *de novo* organ formation in tomato hypocotyl explants. Dark-grown hypocotyl explants (0:24 photoperiod) on sucrose-depleted medium did not produce ARs in the basal region of the explants, and *de novo* organ formation in their apical region was held at stage 1 (**Figure 6e**). On standard photoperiod (16:8) on sucrose-depleted medium, ARs were produced in the basal region of half of the explants, but *de novo* organ formation in the apical region was not observed (**Figure 6e**). To confirm that the observed effect of sucrose on wound-induced organ formation depends on the glycolysis pathway, we grew hypocotyl explants on media supplemented with different sugars during 10 days in the dark (see **Materials and Methods**). When grown in sucrose, fructose or glucose, ARs rapidly emerged and by 5 dae all explants produced at least one AR (**Figure 6f**). Additionally, hypocotyl explants grown on sorbitol-supplemented medium did not produce any AR until the explants were transferred to 16:8 photoperiod at 11 dae (**Figure 6f**). At the end of the experiment (17 dae), we found that hypocotyl explants were able to produce *de novo* shoots (stage 3) with similar proportions (31%-50%) regardless of the sugar that was added (**Figure 6g**). We inhibited glycolysis by incubating hypocotyl explants with 2-deoxyglucose (2-DG) and AR emergence was completely blocked (**Figure 6h, i**). In addition, *de novo* shoot formation was not observed in the apical region of these explants, which were held at stage 1 (**Figure 6i**). These results indicate that exogenous sucrose was essential for effective shoot regeneration in the apical region of the hypocotyl explants, but that was not necessary for AR formation when another energy source is present (e.g. photosynthesis/photorespiration-derived compounds).

## DISCUSSION

We have established a new experimental system using Micro-Tom hypocotyl explants to study wound-induced *de novo* shoot formation and AR development without exogenous application of plant hormones (this work; Larriba et al., in preparation). We employed bulk time-course RNA-seq and target metabolite profiling to characterize tissue-specific reprogramming events in tomato hypocotyl explants. Time-course transcriptome analyses of wound-induced *de novo* organ formation have been performed in *Arabidopsis thaliana* (Chen et al., 2016; Efroni et al., 2016; Ikeuchi et al., 2017; Pan et al., 2019). Besides, transcriptome dynamics during tissue-reunion after wounding have been extensively studied in this species (Asahina et al., 2011; Melnyk et al., 2015; Melnyk et al., 2018). These earlier studies have contributed to the understanding of the hormonal crosstalk driving the different tissue reprogramming programs and have allowed the identification of some of the transcriptional regulators involved, but comparative studies with other plant species are limited. Our transcriptome data indicate both tissue-specific differences and divergent developmental patterns at the apical and basal regions of tomato hypocotyl explants after wounding. We also found significant variation in lncRNA expression across the studied tissue regions and through the time-course, suggesting a regulatory role for these non-protein-coding genes in wound-induced organ formation will be addressed in future studies. In our experimental system, AR formation in the basal region of the hypocotyl relies on the activation of resident stem cells due to quick auxin homeostasis regulation after wounding (Alaguero-Cordovilla et al., 2021; Larriba et al., in preparation), while in the apical region, callus formation and metabolic reprogramming precede *de novo* shoot formation (this work).

We found differential regulation of photosynthesis-related genes in the apical and basal hypocotyl after wounding. On the one hand, photosynthesis-related genes were strongly downregulated in the basal region during the time-course. On the other hand, most of the photosynthesis-related genes were constitutively upregulated in the apical region after wounding, except those encoding antenna proteins (LHC) and PSI components that were transiently downregulated at T1. As these conditions resembled high-light stress adaptation (Johnson and Wientjes, 2020), the high PSII/PSI ratio in the apical region shortly after wounding might lead to the formation of singlet oxygen and thus cause photo-oxidative damage. Indeed, we observed a local increase of ROS production in the apical region at T1, which was downregulated at the later time point, likely by the enhanced expression of genes encoding FD, PRX and TRX, which participate in fine-tuning chloroplast performance under several stress conditions (Cejudo et al., 2020). In agreement with dynamic regulation of photosynthesis function in the apical region after wounding, the expression of key regulators of light acclimation (ABC1K1, STN8 and PSB33) was specifically upregulated in this tissue. DE-ETIOLATED1 (DET1) is a major repressor of photomorphogenesis that associates with CULLIN4 (CUL4)-based E3 ubiquitin ligases for the degradation of several transcription factors (Lau and Deng, 2012). In mammals, Sox2 is a key transcriptional factor for maintaining pluripotency of stem cells and it has been shown that its protein turnover is regulated by the CUL4A^DET1-COP1^ complex (Cui et al., 2018). Whether a conserved regulatory module, downstream of DET1, is required for wound-induced *de novo* shoot formation in tomato hypocotyl explants needs to be addressed. In summary, our results point to blue light as a novel photoregenerative factor in tomato, although the molecular mechanisms that unleash the regenerative signal requires further investigation.

Reprogramming of glucose metabolism in cancer cells even in the presence of atmospheric oxygen (the Warburg effect) is a key event to sustain tumor growth, as high glycolytic flux provides sufficient energy and metabolic intermediates required by the rapidly proliferating cells (Boroughs and Deberardinis, 2015; Yu et al., 2017; Frezza, 2020). Besides, cancer cells rely on the serine/glycine biosynthetic pathway for the production of one-carbon (C1) metabolites required for tumorous growth (Amelio et al., 2014; Li and Ye, 2020). We demonstrated that wound-induced organ formation in tomato hypocotyl explants, both ARs and *de novo* shoot formation, was dependent on the sugar supply. Several sugar transporters of the SWEET family and CWIN were constitutively upregulated in both the apical and the basal regions of the hypocotyl after wounding. Mirroring what has been found during zebrafish heart regeneration (Honkoop et al., 2019), blocking glycolysis with 2-deoxyglucose impaired the ability of cells near the wound to produce new organs. These results suggest that glycolysis-derived energy (ATP) drives cell proliferation in tomato hypocotyl explants required for wound-induced organ formation, and thus these tissues were rapidly reprogrammed as sink tissues as regards sugar assimilation.

We found specific upregulation of the photorespiratory pathway in the apical region of tomato hypocotyl explants after wounding and during *de novo* shoot formation. Photorespiration recycles toxic 2-phosphoglycolate into 3-PGA, which can enter the Calvin–Benson cycle, and it is a key metabolite driving C1 metabolism in plants (Eisenhut et al., 2019). We hypothesized that initial callus growth in the apical region of the hypocotyl relies on photorespiration-produced 3-PGA, which could be used as a substrate for gluconeogenesis too.

Indeed, we found differential upregulation of key genes of gluconeogenesis (*PPDK, FBPase*) that correlated with increased levels of the intermediates of these two pathways (glyoxylate, hydroxypyruvate, 3-PGA, G3P) in the apical region of the explants. By local inhibition of photorespiration in the apical region, we provide additional evidence of a functional role of this pathway to sustain proliferation and callus formation, which preceded and was required for wound-induced shoot initiation. Our results highlight the key role of photorespiration, considered as an energetically inefficient pathway, to trigger wound-induced *de novo* shoot formation, in light with its proposed role to overcome abiotic stresses (Sunil et al. 2019).

Taken together, our results indicate that wounding induced a broad metabolic reprogramming of some cells in the apical region of the hypocotyl (photosynthesis reactivation, photorespiration induction, and high glycolysis), providing energy and structural elements required for rapid proliferation during initial callus growth and that are essential for fate reprogramming required for *de novo* shoot formation.

We found striking patterns of ROS accumulation in the apical and basal region of the hypocotyl during the studied time-course, which was not directly correlated with the extent of cell death (this work), and hence might have a signaling role in *de novo* organ formation. ROS accumulation in the basal region of the hypocotyl explants at T4 and T8 might be related to AR emergence (Mhimdi and Pérez-Pérez, 2020). Indeed, during primary root (PR) and lateral root (LR) development in *Arabidopsis thaliana*, ROS distribution (which is regulated by specific POX enzymes) is required for transition between proliferating and differentiating cells in the meristem (Tsukagoshi et al., 2010; Fernández-Marcos et al., 2017). POX display different functions besides their direct role in wounding response (Minibayeva et al., 2015; Shigeto and Tsutsumi, 2016). Indeed, some of the POX-encoding genes were strongly deregulated in our RNA-seq. Interestingly, a subset of class III POX genes was targeted by LATERAL ORGAN BOUNDARIES DOMAIN29 (LBD29) in Arabidopsis (Xu et al., 2018), a master regulator of auxin-induced callus formation (Fan et al., 2012). Moreover, the LBD29-regulated biological responses largely resemble those of hypoxia treatment, which has been long considered as a crucial factor in determining cell fate in mammals (Xu et al., 2018). We found upregulated in the apical region several *POX* genes (*Prx05*, *Prx11*, *Prx25*, *Prx26*, *Prx38* and *Prx68*) that were structurally related to the LBD targets described above, suggesting that some LBD29-gene regulatory networks might be conserved in tomato. Natural variation in hormone-induced shoot regeneration among Arabidopsis accessions was dependent on the expression of a TRX-encoding gene that directly modulates ROS homeostasis (Zhang et al., 2018). Despite differences in wound response between animals and plants, ROS is a common signal in both systems (Suzuki and Mittler, 2012). In cancer cells, the increase of ROS stabilizes the Hypoxia-Inducible Factor-1α (HIF1α) that upregulates glucose catabolism and allows tumor progression (Nagao et al., 2019). Furthermore, iPSC reprogramming is enhanced under hypoxic conditions through ROS-mediated activation of HIF1α (Hawkins et al., 2016). Importantly, transient ROS signaling during early reprogramming demethylates the promoter of a gene encoding the homeobox protein NANOG required for stem cell pluripotency (Ying et al., 2016). A similar hypoxia-mediated mechanism may be present in plants, but operating on the negative regulators of the ERF-VII transcription factors instead (Pucciariello and Perata, 2021). However, a direct link between ROS production, metabolic reprogramming, and wound-induced organ regeneration in tomato hypocotyl explants awaits experimental confirmation.

## ACKNOWLEDGEMENTS

We thank María José Ñíguez-Gómez for her expert technical assistance.

## ADDITIONAL INFORMATION

### Authors’ contributions

JMPP designed the research. EL, ABSG, CMA and AA performed the research and analyzed the data. JMPP wrote the manuscript with the assistance of EL. All authors read and approved the final manuscript.

### Data Availability Statement

Raw sequence files and read count files are publicly available in the NCBI’s BioProject repository upon publication. Gene functional annotation is available in the supplementary material of this article. All other data that support the findings of this study are available from the corresponding author upon reasonable request.

