## Supplemental Figure for "Tissue-specific metabolic reprogramming during wound induced *de novo* organ formation in tomato hypocotyl explants"

### Supplemental Figure 1

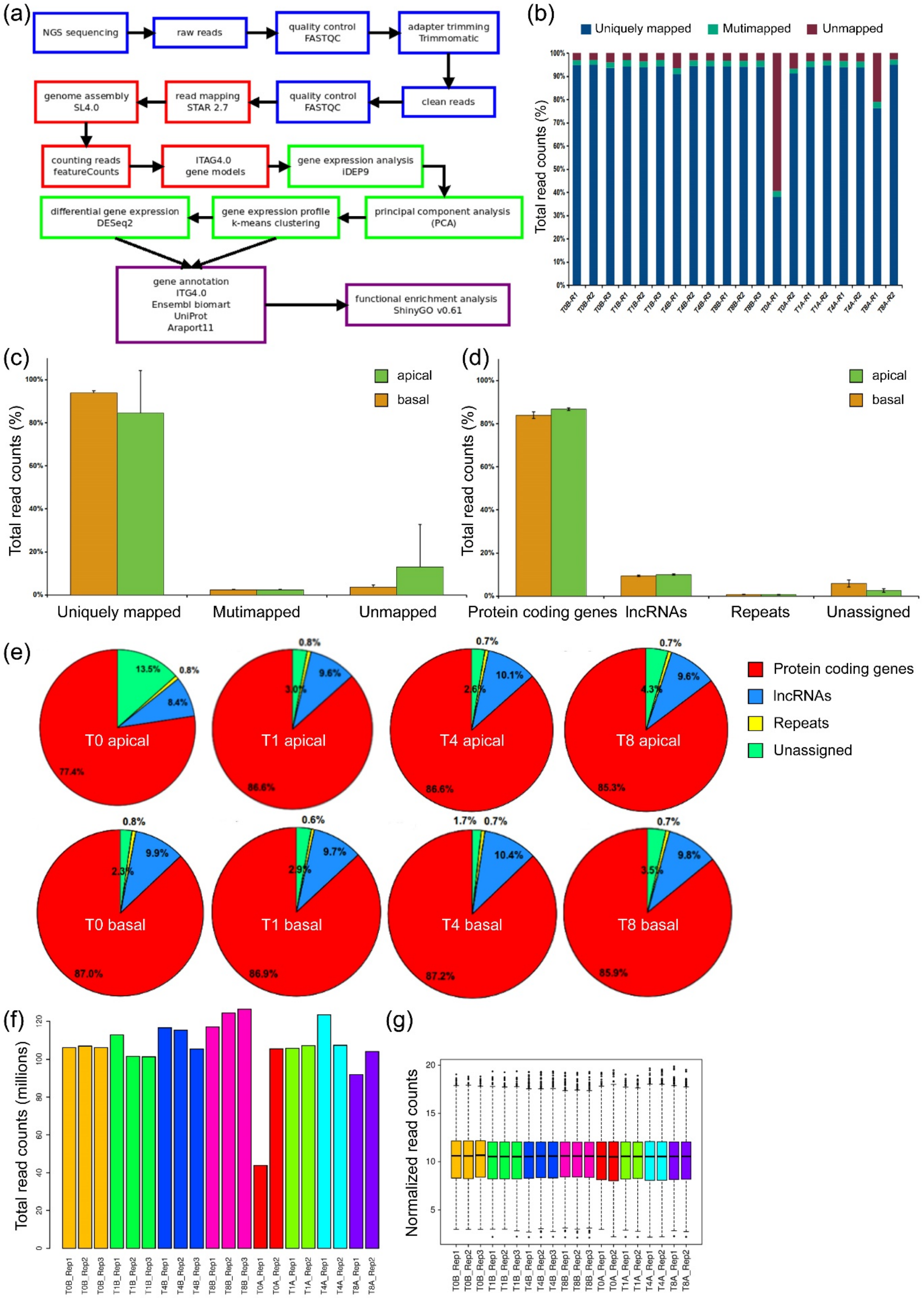


**Supplemental Figure 1**. **Bioinformatics pipeline, read mapping and gene expression statistics.** (a) Bioinformatics workflow. Blue: cleaning and quality evaluation of NGS sequencing libraries; red: mapping and gene counting; green: library normalization and differential expression; purple: functional annotation and GO enrichment analyses. (b) Mapping results of the sequencing libraries. (c) Mapping result summary. (d) Summary of read counts distribution after genome features annotation. (e) Read counts distribution after genome features annotation of the different samples studied. (f) Number of reads assigned to protein coding genes in each sequencing library. (g) Box plot of reads assigned to protein coding genes in each sequencing library after normalization.

#### Supplemental Figure 2

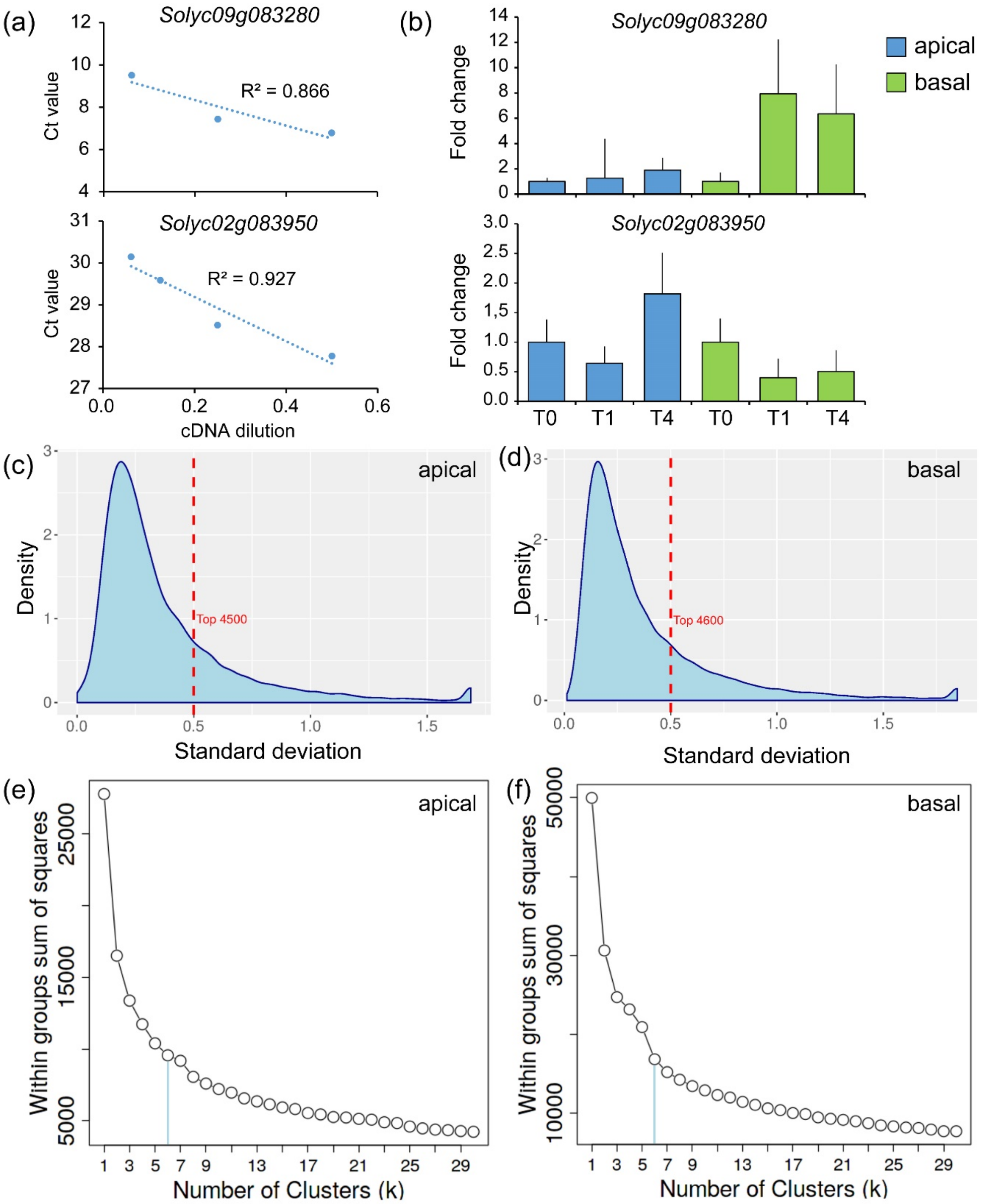


**Supplemental Figure 2. RNA-seq validation by RT-qPCR and K-means clustering analysis.** (a) Primer validation of selected genes. Each dot represents the relative expression data for a given sample. (b) RT-qPCR of the expression of selected genes. Bars indicate normalized expression levels ± standard deviation (SD) relative to the hypocotyl explant at T0. (c, d) Distribution of gene expression SDs of the RNA-Seq results from apical (c) and basal (d) regions of the hypocotyl explants. (e, f) Identification of k-means optimal cluster number by Elbow analysis of the RNA-Seq results from apical (e) and basal (f) regions of the hypocotyl explants.

#### Supplemental Figure 3

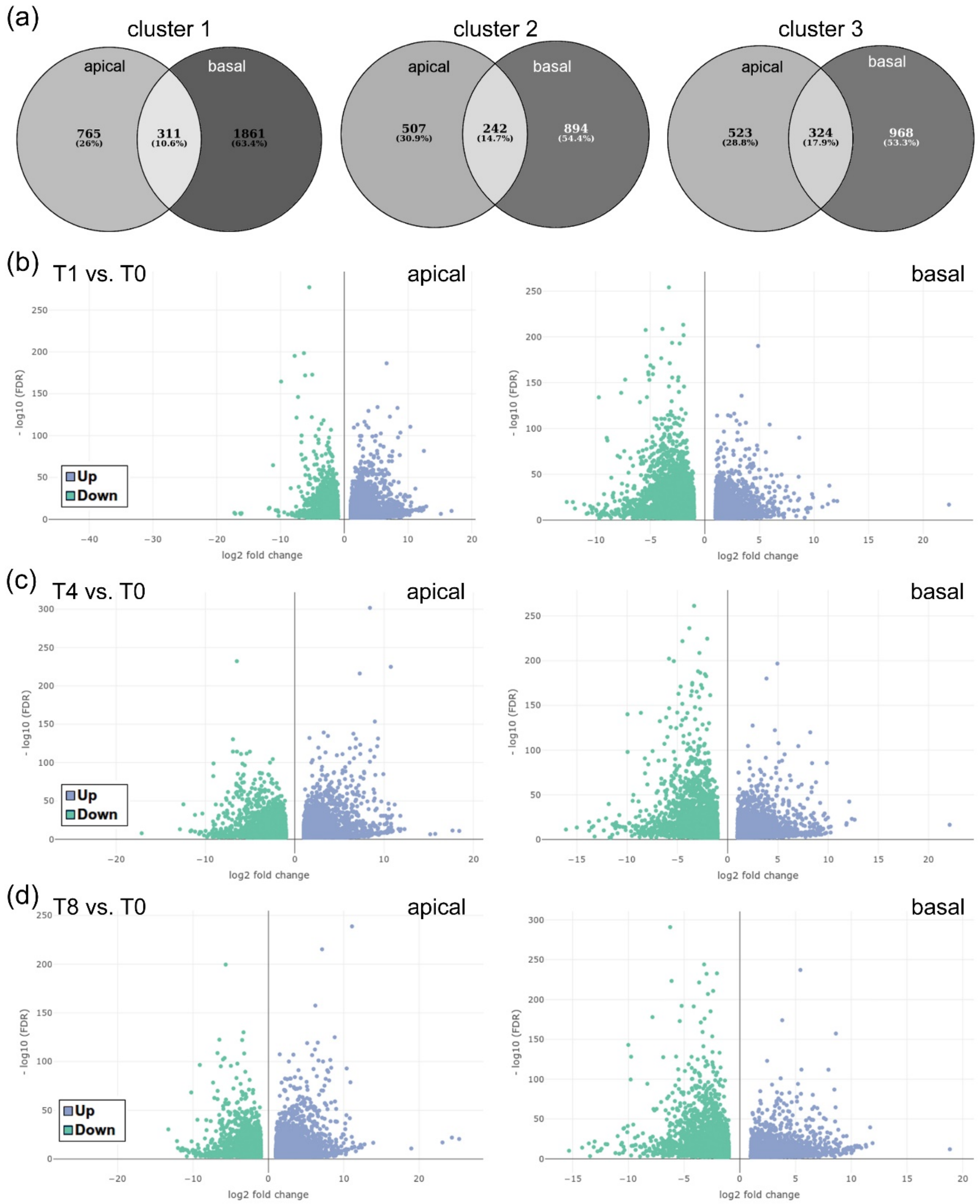


**Supplemental Figure 3. Venn diagrams and volcano plots of the RNA-seq results.** (a) Venn diagrams from shared k-means clusters between the apical and basal regions of the hypocotyl explants. (b-d) Volcano plots of contrast between T1 and T0 (a), T4 and T0 (c) and T8 and T0 (d) from apical and basal regions of the hypocotyl explants. The $\log_2$fold change is plotted on the x-axis, and the negative $\log_{10}$ (FDR) is plotted on the y-axis. The green/blue dots indicate the differentially expressed genes (DEG) with the absolute value of $\log_2$fold change> < |1| and FDR<0.01.

### Supplemental Figure 4

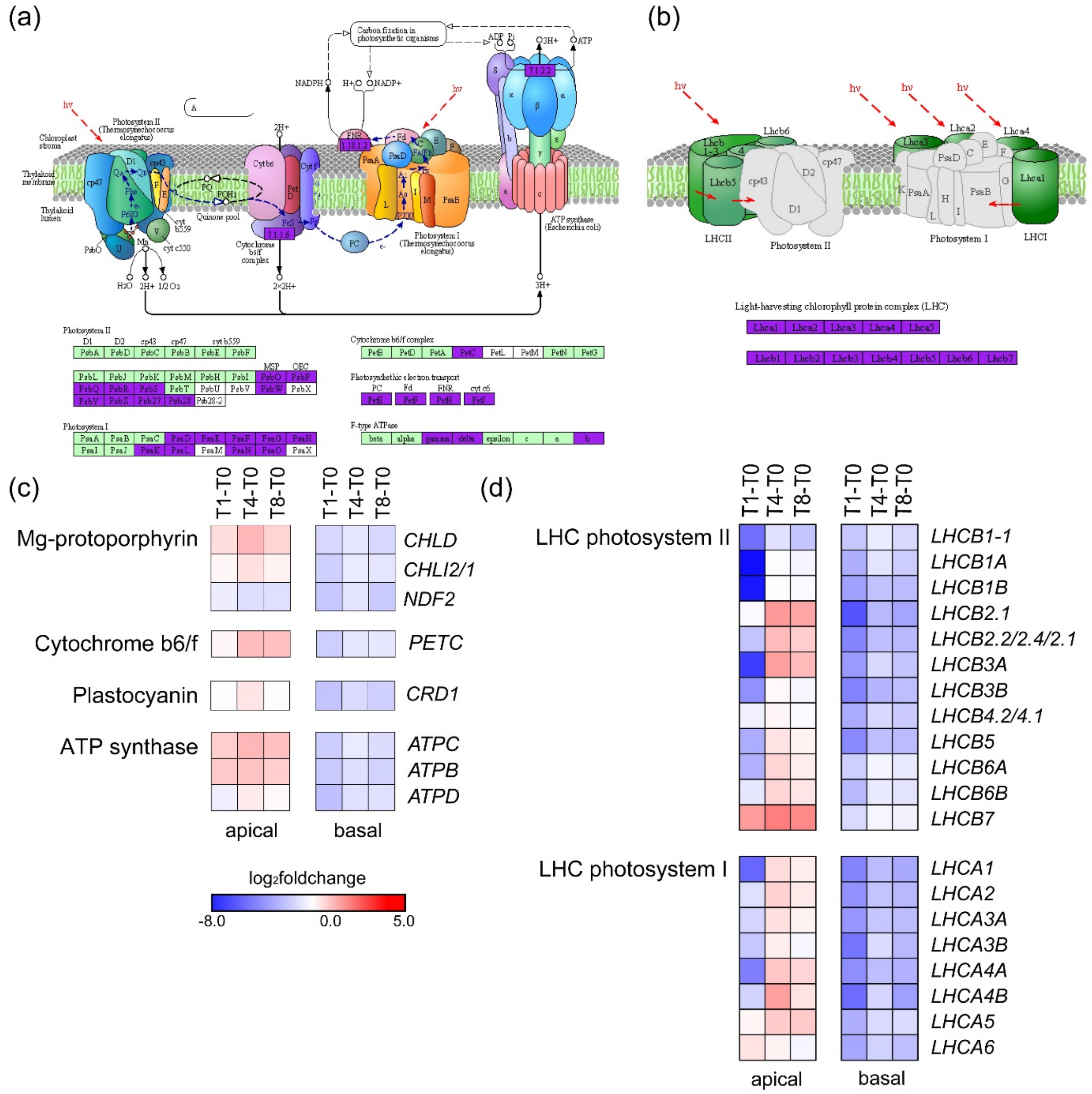


**Supplemental Figure 4. Expression of photosynthesis-related genes during wound-induced organ formation.** (a) Photosynthesis pathway and (b) light-harvesting chlorophyll protein complexes (LHC) based on the Kyoto Encyclopedia of Genes and Genomes (KEGG) database (Kanehisa et al., 2021); *hv* designates light energy/photons. Genes expressed in our RNA-seq are indicated in light green, and those deregulated were indicated in purple. (c, d) DEG involved in know photosynthesis subcomplexes (c), or encoding the antenna proteins of the LHC (d). Gene annotations are found in Supplemental Table 4.

#### Supplemental Figure 5

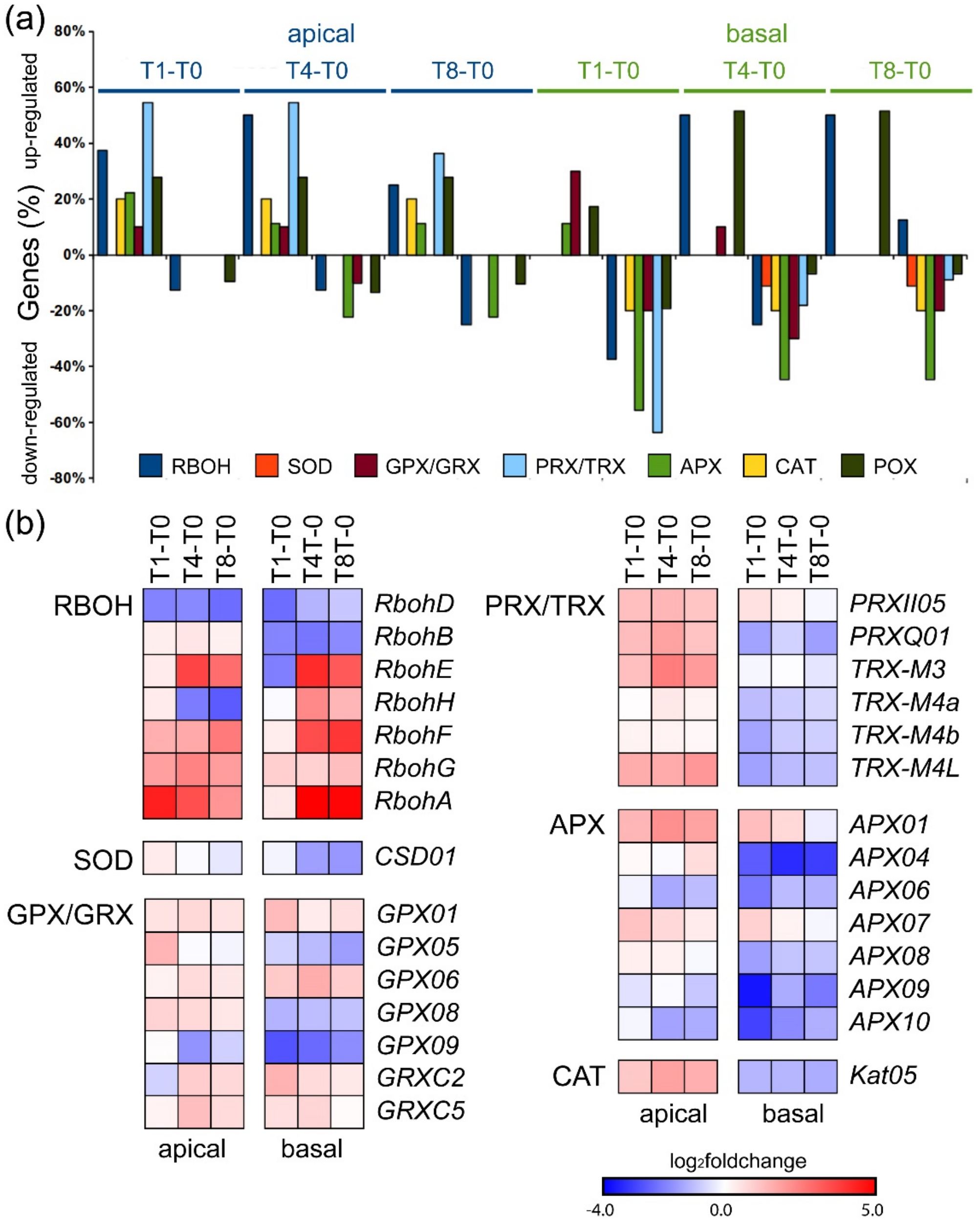


**Supplemental Figure 5. Expression of ROS related genes during wound-induced organ formation.** (a) Percentage of deregulated genes of the studied oxidoreductases families in the apical and basar region of the hypocotyl. (b) DEG encoding ROS production and scavenging enzyme. RBOH, NADPH oxidase/respiratory burst oxidase homolog; SOD, superoxide dismutase; POX, peroxidase; GPX, glutathione peroxidase; PRX/TRX, peroxiredoxins/thioredoxins; APX, ascorbate peroxidases; CAT, catalases. Gene annotations are found in Supplemental Table 5.

### Supplemental Figure 6

(a)

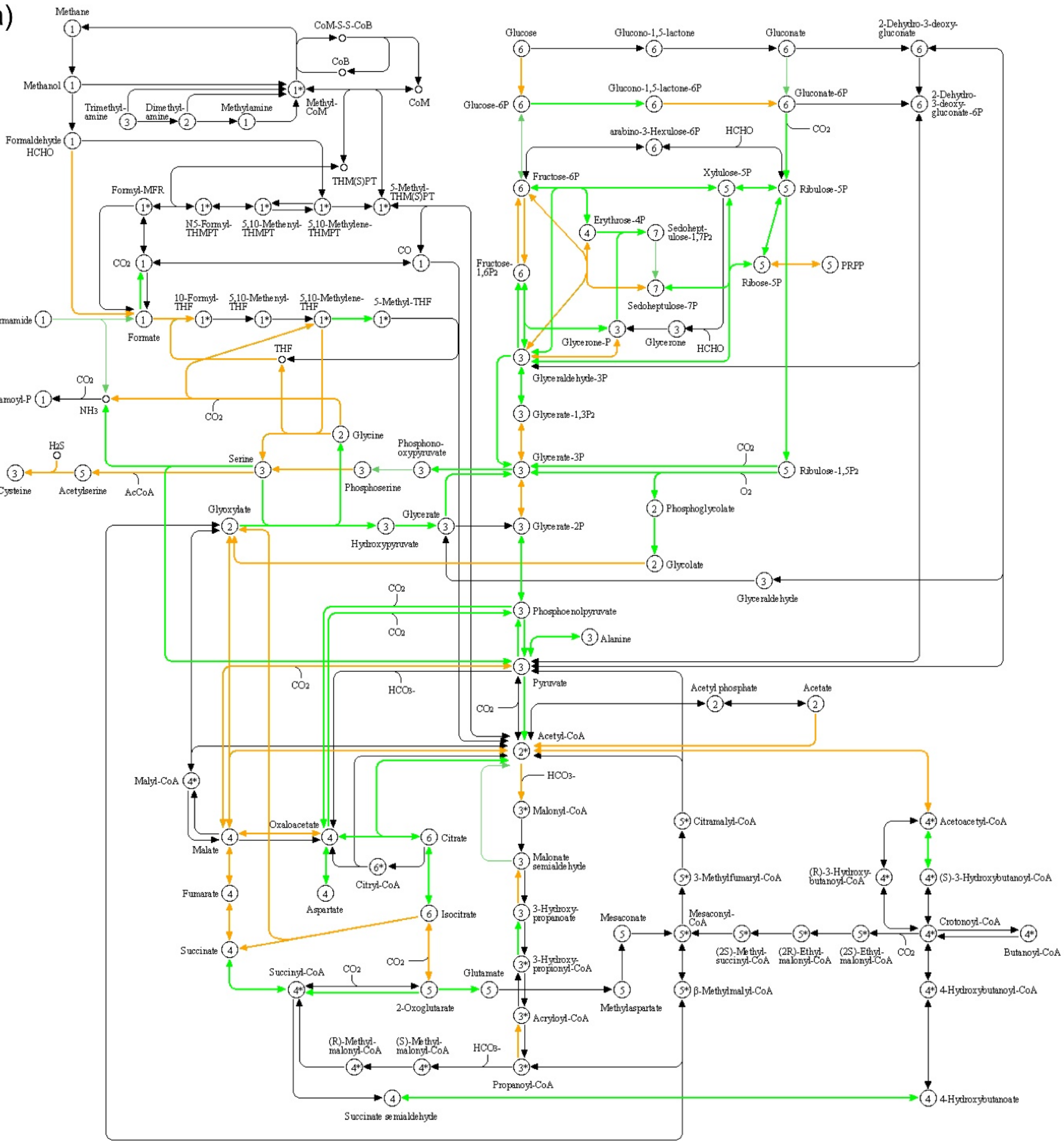

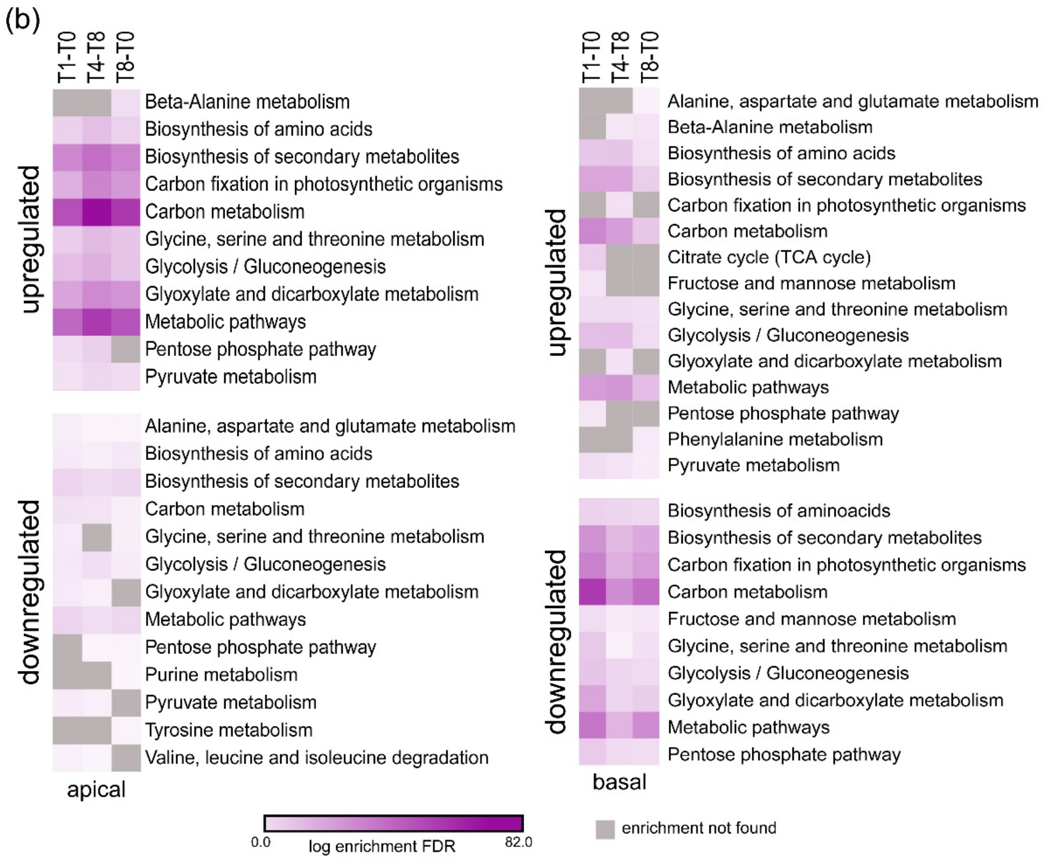


**Supplemental Figure 6. Deregulation of carbon metabolism genes during wound-induced organ formation.** (a) Carbon metabolism pathway based on the KEGG database (Kanehisa et al., 2021). Green and orange arrows respectively represent genes found deregulated in at least one of the contrast, and genes expressed but not found deregulated in our RNA-seq. (b) Enrichment of specific carbon metabolism modules. Gene annotations are found in Supplemental Table 6.

### SUPPLEMENTAL TABLES

**Supplemental Table 1. Primers for RT-qPCR validation of RNA-seq data.** Gene name and gene identifier in ITAG4.0 annotation (Solyc ID) are included. Nucleotide sequence of the forward and reverse primers and amplicon size (cDNA) are indicated.

**Supplemental Table 2. Biological process (BP) GO term enrichment analysis from k-means gene clustering.** List of BP GO terms found in enrichment analysis of specific and common genes found in k-means clustering. Common genes of apical and basal tissues, as well as the specific expression genes of both tissues, for clusters C1, C2 and C3 (see Figure 1) were indicated as: exclusive apical, exclusive basal and common, respectively. In addition, the enriched terms belonging to clusters 4 and 5, specific to the apical tissue, are shown as clusters 4 and 5, respectively. For each group of genes, the FDR value obtained in the enrichment analysis is indicated, the genes present in each analyzed group belonging to that term, as well as the number of genes noted in the background for that term. BP functional terms were sorted according to fold enrichment and FDR<0.01.

**Supplemental Table 3. Biological process (BP) GO term enrichment analysis from DEG.** In each Venn diagram subset we compared upregulated and downregulated DEG in the apical and basal regions from three different contrast, T1-T0, T4-T0 and T8-T0 (see Figure 2d, e). Functional categories are sorted according to fold enrichment and FDR<0.01.

**Supplemental Table 4. Annotation of photosynthesis-related genes in tomato.** SolycID and ITAG4.0 annotation were retrieved from SolGenomics (https://solgenomics.net/). Putative *Arabidopsis thaliana* orthologs were identified from the Ensembl Plants database using BioMart. . %id. Target A.thaliana, percentage of identity of target *Arabidopsis thaliana* gene identical to tomato gene; %id. tomato gene, percentage of identity of target tomato gene identical to *Arabidopsis thaliana* gene; confidence, orthology confidence score from the Emsembl Plants database. KEGG identifiers and annotations were retrieved using a strategy RBBH as described in M&M section.

**Supplemental Table 5. Annotation and classification of genes associated to reactive oxygen species (ROS) production and detoxification.** Gene annotation was retrieved from indicated sources.

**Supplemental Table 6. Annotation of carbon metabolism genes in tomato.** See Supplemental Table 4 legend for details.

**Supplemental Table 7. Annotation of genes encoding sugar transporters, invertases and sucrose syntheses in tomato.** Gene annotation was retrieved from the indicated source.
